# Cellular responses to reactive oxygen species can be predicted on multiple biological scales from molecular mechanisms

**DOI:** 10.1101/227892

**Authors:** Laurence Yang, Nathan Mih, Amitesh Anand, Joon Ho Park, Justin Tan, James T. Yurkovich, Jonathan M. Monk, Colton J. Lloyd, Troy E. Sandberg, Sang Woo Seo, Donghyuk Kim, Anand V. Sastry, Patrick Phaneuf, Ye Gao, Jared T. Broddrick, Ke Chen, David Heckmann, Richard Szubin, Ying Hefner, Adam M. Feist, Bernhard O. Palsson

## Abstract

Catalysis using iron-sulfur clusters and transition metals can be traced back to the last universal common ancestor. The damage to metalloproteins caused by reactive oxygen species (ROS) can completely inhibit cell growth when unmanaged and thus elicits an essential stress response that is universal and fundamental in biology. We develop a computable multi-scale description of the ROS stress response in *Escherichia coli*. We show that this quantitative framework allows for the understanding and prediction of ROS stress responses at three levels: 1) pathways: amino acid auxotrophies, 2) networks: the systemic response to ROS stress, and 3) genetic basis: adaptation to ROS stress during laboratory evolution. These results show that we can now develop fundamental and quantitative genotype-phenotype relationships for stress responses on a genome-wide basis.

---

All aerobic life requires management of corrosive oxidative stress. Oxygen toxicity is manifested in damage to cellular components by reactive oxygen species (ROS). Cells generate ROS endogenously when flavin, quinol, or iron cofactors are autoxidized ^1^. ROS species are known to damage DNA, certain iron-containing metalloproteins, and other cellular processes ^2^. In addition to inherent endogenous ROS production, eukaryotes and microbes produce exogenous ROS as H2O2, superoxide, or redox-cycling compounds to inflict oxidative stress on competitors ^3^. In spite of the fundamental importance of ROS damage on cellular functions, we lack a framework that connects known and hypothesized individual molecular targets of ROS to systemic physiological responses. Here, we address this gap using a genome-scale computational systems biology approach focused on the processes that determine homeostasis of iron, which is essential for *E. colf’s* growth, yet is vulnerable to ROS.

We developed a computable multi-scale description of ROS damage to metalloproteins in the context of metabolic function and macromolecular expression in *Escherichia coli* MG1655 ^4–8^. To mechanistically account for metalloprotein inactivation by ROS, we integrated metal cofactor damage and repair pathways for 43 protein complexes involving mononuclear iron or iron-sulfur clusters (Fig. 1). While all of these enzymes are potentially inactivated, the extent of ROS damage has not been experimentally determined for most of them. Certain iron metalloproteins have been shown to avoid inactivation by ROS due to structural properties including full incorporation or solvent accessibility ^9^. To account for these cases, we integrated protein structural property calculations ^10,11^ into a Bayesian network to compute the probability of cofactor inactivation by ROS (Fig. S1). The resulting description is a computable model, referred to as OxidizeME, that accounts for 1,581 proteins that collectively represent up to 85% of the proteome by mass (Data Table S1). OxidizeME makes clear interpretations and predictions of ROS responses, three of which are addressed below; 1) ROS and amino acid auxotrophies, 2) the systemic response to ROS stress, and 3) adaptation to ROS stress during laboratory evolution.

**Fig. 1.**
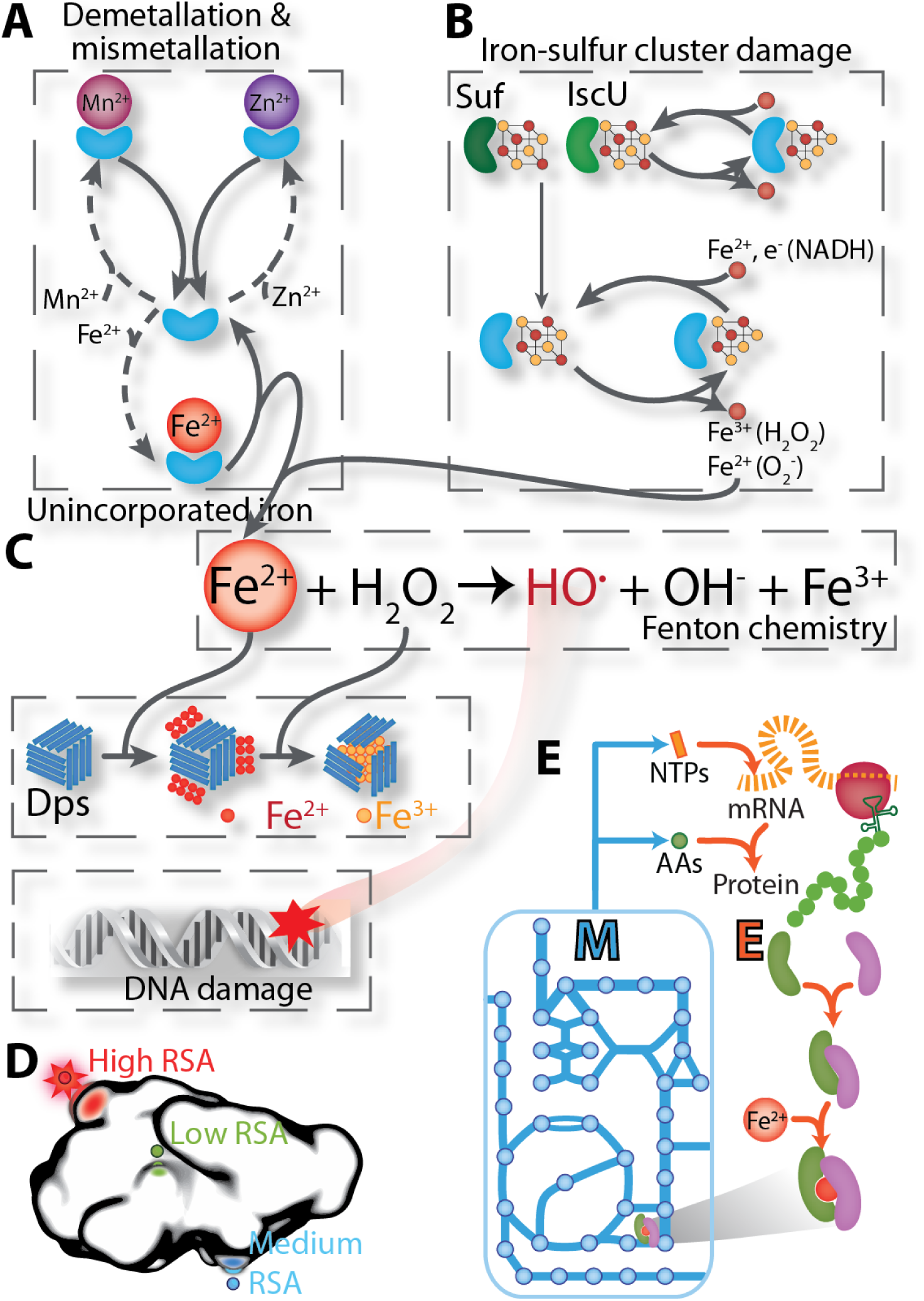
OxidizeME: a multi-scale description of <u>m</u>etabolism and macromolecular <u>e</u>xpression that accounts for damage by reactive oxygen species (ROS) to macromolecules. **(A)** Mononuclear Fe(II) proteins are demetallated by ROS and mismetallated with alternative divalent metal ions. **(B)** Iron-sulfur clusters are oxidized and repaired. **(C)** Unincorporated Fe(II) spontaneously reacts with H_2_O_2_ via Fenton chemistry, generating hydroxyl radicals that damage DNA, while the Dps protein stores unincorporated iron and protects DNA from damage. **(D)** Protein structural properties are computed to estimate the probability of metal cofactor damage by ROS (RSA: relative solvent accessibility). **(E)** Processes in A-D are integrated into a multi-scale oxidative model, named OxidizeME. OxidizeME is used to compute the scope of macromolecular damage and the cellular response for varying intracellular concentrations of superoxide, hydrogen peroxide, and divalent metal ions (Fe(II), Mn(II), Co(II), Zn(II)), See SI for details.

A hallmark response to ROS damage for *E. coli* is the deactivation of branched-chain and aromatic amino acid biosynthesis pathways, which is alleviated by supplementing these amino acids (2). Compared to supplementing all 20 amino acids, OxidizeME correctly predicted that excluding Ile and Val had a greater impact on growth rate than did excluding Phe, Trp, and Tyr (Fig. 2A). The reason that *E. coli* cannot grow under ROS stress without supplementation of branched-chain amino acids is that the iron-sulfur clusters of dihydroxy-acid dehydratase and isopropylmalate isomerase are inactivated by ROS, thus debilitating the branched-chain amino acid biosynthetic pathway ^12^. The auxotrophy for aromatic amino acids was originally attributed to inactivation of the transketolase reaction ^13^, but was recently traced to the mismetallation of the mononuclear iron cofactor in 3-deoxy-D-arabinheptulosonate 7-phosphate (DAHP) synthase ^14^. OxidizeME correctly predicted these molecular mechanisms and their phenotypic consequences (Fig. 2).

**Fig. 2.**
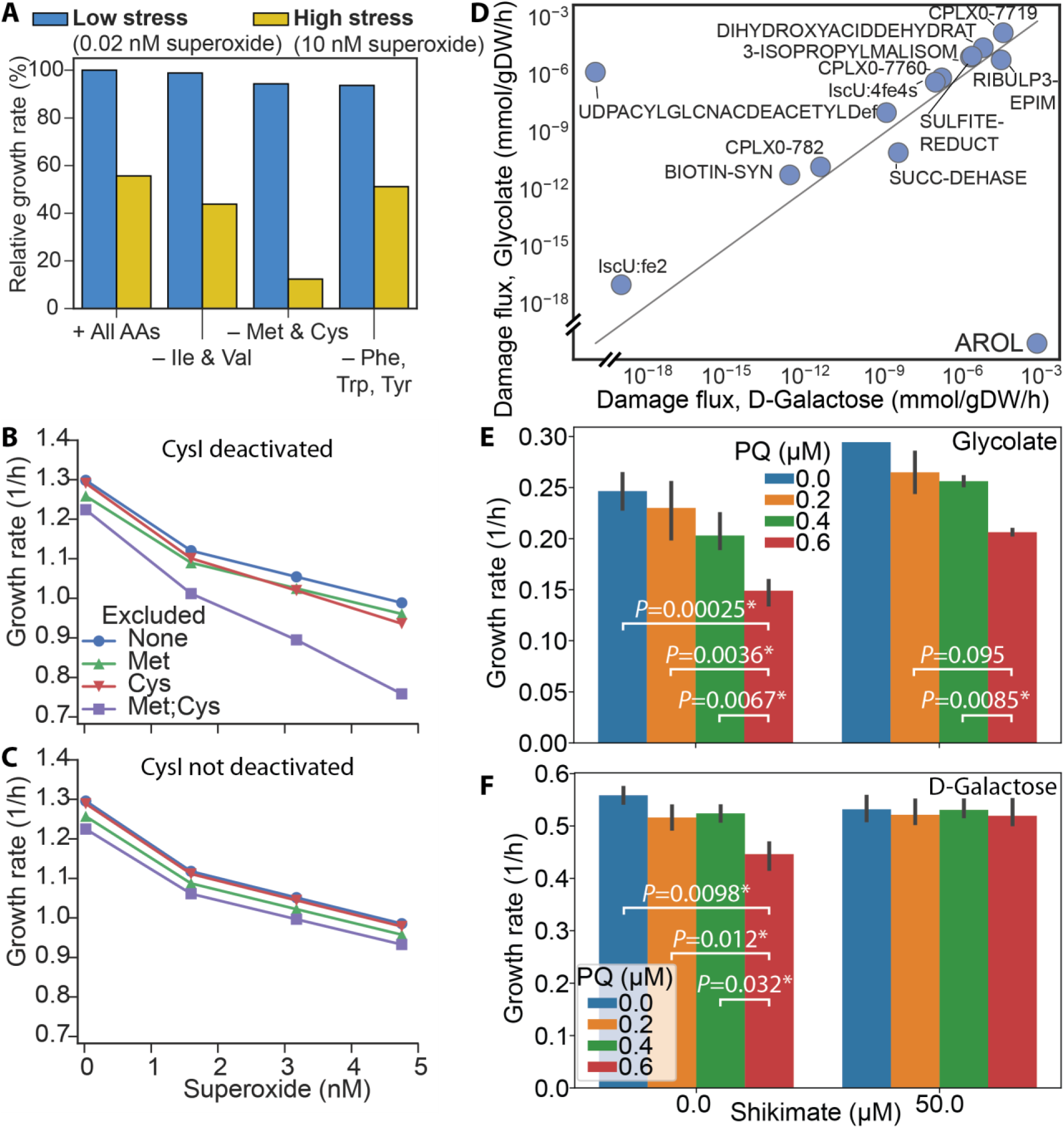
Systemic consequences of ROS stress. **(A)** Predicted growth rate under low and high superoxide concentrations with different supplementation of amino acids (AAs). All AAs refers to all 20 common amino acids, and “-Ile & Val” means all amino acids except Ile and Val were supplemented. **(B)** Predicted growth rate versus superoxide concentration in various sulfurous amino acid supplementation media. **(C)** Same as **(B)** but simulated without damage to CysI by ROS. **(D)** Simulated damage fluxes for growth on glycolate versus D-galactose. AROL: shikimate kinase II. **(E)** Growth rate of MG1655 on glycolate minimal medium with 0 to 0.6 μM PQ, with and without 50 μM shikimate supplementation. **(F)** Same as (E) but for growth on D-galactose minimal medium. * denotes that the growth rate changes significantly between two pQ concentrations (two-tailed Welch’s t-test, *P*<0.01).

Meanwhile, the basis of sulfurous amino acid auxotrophy in *E. coli* remains inconclusive despite multiple investigations ^15,16^. OxidizeME correctly predicted auxotrophy for sulfurous amino acids (cysteine and methionine) under ROS stress (Fig. 2A). We traced a plausible mechanism to damage of the iron-sulfur cluster in CysI, which catalyzes the sulfite reductase step of Cys biosynthesis. Sulfite reductase binds four cofactors: iron-sulfur, FAD, FMN, and siroheme. Consistent with prior studies ^17^, our structural model estimated the siroheme group of sulfite reductase to be difficult to reach by ROS, mainly due to the depth of the cofactor binding residue (Data Table S2). Previous studies showed that the iron-sulfur cluster is likely not autoxidized with molecular oxygen because it is not solvent-exposed ^17^. However, our structural model predicted that the iron-sulfur cluster is reached by ROS when considering both solvent exposure and depth of the cluster-binding residue from the solvent accessible surface (Data Table S2). Simulations confirmed that alleviating damage to sulfite reductase was sufficient to reverse the observed growth rate defect and enable growth at higher ROS concentrations in the absence of Cys and Met (Fig. 2B,C). Our hypothesis that sulfite reductase is deactivated by ROS is consistent with studies in *Salmonella enterica* showing that the activity of this enzyme is indeed reduced by elevated superoxide ^18^. Furthermore, the deactivation of sulfite reductase is consistent with accumulation of its substrate, sulfite, and explains the previously-observed accumulation of sulfite ^16^. We note that CysI inactivation does not exclude the possibility that superoxide additionally leads to cell envelope damage, facilitating leakage of small molecules ^19^. Thus, OxidizeME can be used to understand and predict the basis for amino acid auxotrophies as a systemic response to specific macromolecular vulnerabilities to ROS.

To investigate how environmental context affects ROS tolerance, we simulated growth under superoxide stress in 180 carbon sources (Fig. S2). We then compared pairs of carbon sources in terms of the complexes that are most damaged by ROS. In particular, from simulations we predicted that a key bottleneck to growth on D-galactose under ROS stress is inactivation of shikimate kinase II, AroL (Fig. 2D). In contrast, AroL was predicted to not be a direct bottleneck to growth on glycolate (Fig. 2D). To validate this prediction, we measured growth of *E. coli* MG1655 on these two carbon sources in 0 to 0.6 μΜ paraquat (PQ). PQ is a divalent cation that is taken up opportunistically, typically by polyamine transmembrane transporters, and then undergoes reduction and autoxidation cycles catalyzed by any of three *E. coli* PQ diaphorases to generate superoxide ^20^. To directly test whether AroL is a bottleneck, we also supplemented the cultures with 50 μΜ shikimate. As predicted, shikimate did not alleviate PQ-induced growth defects during growth on glycolate (Fig. 2E). Meanwhile, shikimate alleviated growth defects by PQ during growth on D-galactose (Fig. 2F). These results confirm that OxidizeME is able to accurately predict cell phenotype under ROS stress and this predicted capability is rooted in its ability to compute molecular and macromolecular mechanisms.

Next, we assessed the systemic response of *E. coli* to ROS stress. We measured the transcriptome of *E. coli* under superoxide stress using PQ treatment and identified 914 differentially expressed genes (DEGs), of which 501 were accounted for in OxidizeME (Fig. 3, Fig. S3). In particular, 87 genes were up-regulated. Using OxidizeME, we determined that these 87 genes were more likely activated due to damage that is specific to iron metalloproteins than to any other protein (P<0.001) (see Supplementary Text). Furthermore, of the DEGs that were correctly predicted, a large fraction (84%) of the repressed genes changed due to decreased growth rate from PQ treatment, while 95% of the activated genes were specific responses to stress (Fig. 3). Gene expression is expected to respond to ROS stress directly--e.g., by up-regulating ROS detoxification genes--and indirectly--in response to decreased metabolic rates caused by ROS damage. The responses we identified as being specific to ROS, not growth rate, spanned eight cellular processes (Fig. 3): ROS detoxification, central metabolism, anaerobic respiration, amino acid biosynthesis, cofactor synthesis and repair, translation, iron homeostasis, and transcriptional regulation by the *rpoS* sigma factor.

**Fig. 3.**
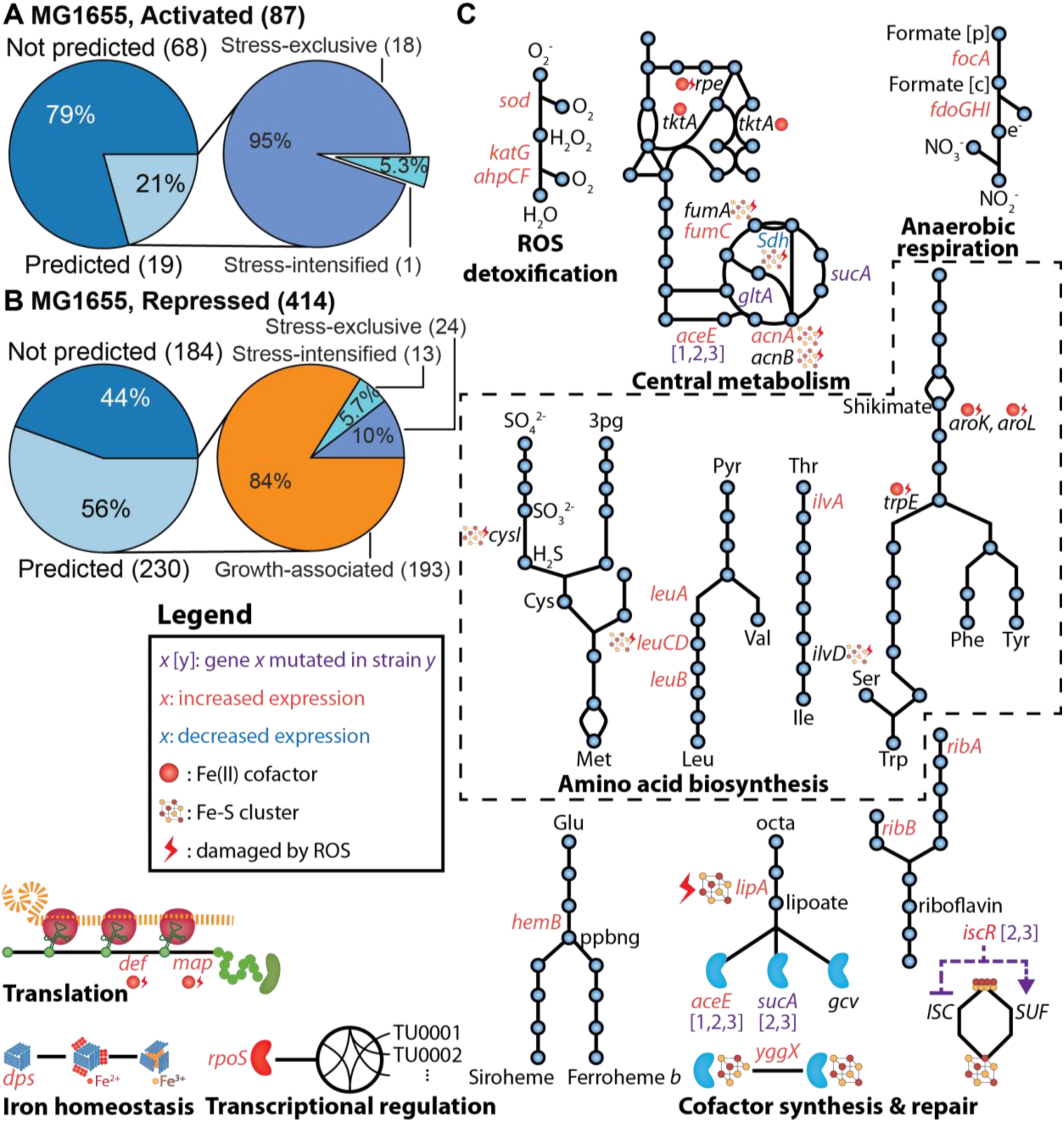
Validation of the consequences and responses to ROS stress. Differentially expressed genes (DEGs) (|log_2_(fold-change)|>0.9|, *FDR*<0.01) that are activated **(A)** and repressed **(B)**. Correctly predicted DEGs are distinguished from global growth-associated regulation using OxidizeME. **(C)** Cellular processes involved in a systemic response to iron metalloprotein damage by ROS.

To investigate the genetic basis for ROS tolerization, we deployed laboratory evolution to investigate how microbes adapt to tolerate high ROS stress. We adaptively evolved a glucose-optimized strain of *E. coli* ^21^, named GLU, in parallel cultures to tolerate three different concentrations of PQ (400, 600, and 800 μM). Re-sequencing identified a common set of genetic changes consisting of mutations in *ygfZ* (folate-binding protein), *aceE* (pyruvate dehydrogenase), and at least one gene in the citric acid cycle: *sucA* (alpha-ketoglutarate dehydrogenase), *gltA* (citrate synthase), or *icd* (isocitrate dehydrogenase) (Table S1). The two most ROS-tolerant strains additionally had mutations in *iscR* (iron-sulfur cluster regulator), while *arnA* (bifunctional, UDP-4-amino-4-deoxy-L-arabinose formyltransferase/UDP-glucuronate dehydrogenase) or *pitA* (metal phosphate:H+ symporter) were mutated in each of these two strains. These results revealed that relatively few mutations confer microbial ROS tolerance, unlike adaptation to thermal stress, which can involve dozens to hundreds of mutations ^22^.

We performed RNA-Seq measurements on the PQ-adapted strains and found cellular responses that contrasted with those of the wild-type (MG1655) and with GLU. These differences also contrast with the conventional understanding of the ROS response. ROS damages components of the iron-sulfur cluster (ISC) assembly pathways through mismetallation of labile iron-sulfur clusters on the scaffold proteins IscU and SufA ^23^. Under ROS stress, conventional understanding would suggest that the ISC assembly pathway is repressed and the sulfur assimilation (SUF) system is up-regulated ^2,24–26^.

IscR regulates the transcription of both ISC and SUF based on coordination of 2Fe-2S at its Cys92, Cys98, and Cys104 residues ^24,27^. In the two most ROS-tolerant strains, we observed the mutation C104S in *iscR*. This mutation may hinder IscR’s ability to incorporate 2Fe-2S and therefore to regulate expression of the ISC and SUF systems under ROS stress. Consistent with deactivation of IscR, we observed expression of ISC and SUF in PQ-adapted strains that was opposite to the unadapted response, where ISC genes are repressed and SUF genes are up-regulated. When treated with 0.6 mM PQ, the most ROS-tolerant strain (PQ3) down-regulated the entire *sufABCDSE* transcription unit (Data Table S3). Furthermore, compared to GLU, the PQ3 strain showed higher expression of the ISC pathway by up-regulating the *iscRSUA* and *hscBA-fdx-iscX* transcription units in 0.2 mM PQ (Data Table S3).

This evolved response directly opposes that of wild-type MG1655 and GLU, which both down-regulated ISC genes and up-regulated *sufA* (Data Table S3). These results show that *E. coli* does not necessarily rely on SUF under ROS stress, and that up-regulation of ISC may be integral for adaptation to higher ROS concentrations. OxidizeME correctly predicted up-regulation of the ISC system and repression of *sufABCDSE* under high ROS stress based on the relative cost-efficiency of Fe-S cluster biosynthesis from each respective system (Data Table S4). Additional observed genetic and expression differences in ROS response between ROS-evolved and unevolved strains are described in Supplementary Text.

The use of iron-sulfur clusters and transition metals to catalyze biological processes can be traced back to the last universal common ancestor ^28^ and ROS stress has a profound impact on all aerobic life forms. Thus, OxidizeME represents a fundamental advance in our understanding of stress-response mechanisms as it provides a genome-wide description of metabolism, protein expression, prosthetic group engraftment, and ROS protecting mechanisms, that all together account for up to 85% of the proteome by mass. Finally, the ability to quantitatively and mechanistically describe responses to ROS damage of ancient conserved molecular targets has broad implications for organisms across the tree of life.

## Acknowledgments

This work was supported by the National Institute of General Medical Sciences of the National Institutes of Health Grants U01 GM12098 and R01 GM057089, and Novo Nordisk Foundation Grant NNF10CC1016517. This research used resources of the National Energy Research Scientific Computing Center, a DOE Office of Science User Facility supported by the Office of Science of the U.S. Department of Energy under Contract No. DE-AC02-05CH11231. All data reported in the paper are available in the Supplementary Materials.

## Supplementary Materials

Materials and Methods

Supplementary Text

Figures S1-S4

Table S1

Data Tables S1-S4

References (*29-47*)

## References and Notes

1. Brynildsen, M. P., Winkler, J. a, Spina, C. S., MacDonald, I. C. & Collins, J. J. Potentiating antibacterial activity by predictably enhancing endogenous microbial ROS production. Nat. Biotechnol. 31, 160–165 (2013).

2. Imlay, J. a. The molecular mechanisms and physiological consequences of oxidative stress: lessons from a model bacterium. Nat. Rev. Microbiol. 11, 443–54 (2013).

3. Imlay, J. A. Iron-sulphur clusters and the problem with oxygen. Mol. Microbiol. 59, 1073–1082 (2006).

4. Thiele, I. et al. Multiscale modeling of metabolism and macromolecular synthesis in E. coli and its application to the evolution of codon usage. PLoS One 7, e45635 (2012).

5. O’Brien, E. J., Lerman, J. A., Chang, R. L., Hyduke, D. R. & Palsson, B. Ø. Genome-scale models of metabolism and gene expression extend and refine growth phenotype prediction. Mol. Syst. Biol. 9, 693 (2013).

6. Lloyd, C. J. et al. COBRAme: A Computational Framework for Models of Metabolism and Gene Expression. (2017). doi: 10.1101/106559

7. Chen, K. et al. Thermosensitivity of growth is determined by chaperone-mediated proteome reallocation. Proc. Natl. Acad. Sci. U. S. A. 114, 11548–11553 (2017).

8. Yang, L., Yurkovich, J. T., King, Z. A. & Palsson, B. O. Modeling the multi-scale mechanisms of macromolecular resource allocation. Curr. Opin. Microbiol. 45, 8–15 (2018).

9. Sobota, J. M. & Imlay, J. a. Iron enzyme ribulose-5-phosphate 3-epimerase in Escherichia coli is rapidly damaged by hydrogen peroxide but can be protected by manganese. Proc. Natl. Acad. Sci. U. S. A. 108, 5402–5407 (2011).

10. Brunk, E. et al. Systems biology of the structural proteome. BMC Syst. Biol. 10, (2016).

11. Mih, N. et al. ssbio: A Python Framework for Structural Systems Biology. Bioinformatics (2018). doi:10.1093/bioinformatics/bty077

12. Macomber, L. & Imlay, J. A. The iron-sulfur clusters of dehydratases are primary intracellular targets of copper toxicity. Proc. Natl. Acad. Sci. U. S. A. 106, 8344–9 (2009).

13. Benov, L. & Fridovich, I. Why superoxide imposes an aromatic amino acid auxotrophy on Escherichia coli: The transketolase connection. J. Biol. Chem. 274, 4202–4206 (1999).

14. Sobota, J. M., Gu, M. & Imlay, J. a. Intracellular hydrogen peroxide and superoxide poison 3-Deoxy-D-Arabinoheptulosonate 7-phosphate synthase, the first committed enzyme in the aromatic biosynthetic pathway of Escherichia coli. J. Bacteriol. 196, 1980–1991 (2014).

15. Pollak, N., Dölle, C. & Ziegler, M. The power to reduce: pyridine nucleotides--small molecules with a multitude of functions. Biochem. J. 402, 205–18 (2007).

16. Benov, L. & Fridovich, I. Superoxide Imposes Leakage of Sulfite fromEscherichia coli. Arch. Biochem. Biophys. 347, 271–274 (1997).

17. Messner, K. R. & Imlay, J. A. The identification of primary sites of superoxide and hydrogen peroxide formation in the aerobic respiratory chain and sulfite reductase complex of Escherichia coli. J. Biol. Chem. 274, 10119–28 (1999).

18. Thorgersen, M. P. & Downs, D. M. Oxidative stress and disruption of labile iron generate specific auxotrophic requirements in Salmonella enterica. Microbiology 155, 295–304 (2009).

19. Benov, L., Kredich, N. M. & Fridovich, I. The mechanism of the auxotrophy for sulfur-containing amino acids imposed upon Escherichia coli by superoxide. J. Biol. Chem. 271, 21037–40 (1996).

20. Liochev, S. I. & Fridovich, I. Paraquat diaphorases in Escherichia coli. Free Radic. Biol. Med. 16, 555–559 (1994).

21. LaCroix, R. A. et al. Use of adaptive laboratory evolution to discover key mutations enabling rapid growth of Escherichia coli K-12 MG1655 on glucose minimal medium. Appl. Environ. Microbiol. 81, 17–30 (2015).

22. Chang, R. L. et al. Structural systems biology evaluation of metabolic thermotolerance in Escherichia coli. Science 340, 1220–3 (2013).

23. Ranquet, C., Ollagnier-de-Choudens, S., Loiseau, L., Barras, F. & Fontecave, M. Cobalt stress in Escherichia coli. The effect on the iron-sulfur proteins. J. Biol. Chem. 282, 30442–51 (2007).

24. Mettert, E. L. & Kiley, P. J. Coordinate regulation of the Suf and Isc Fe-S cluster biogenesis pathways by IscR is essential for viability of Escherichia coli. J. Bacteriol. 196, 4315–23 (2014).

25. Lill, R. Function and biogenesis of iron–sulphur proteins. Nature 460, 831–838 (2009).

26. Lee, J.-H., Yeo, W.-S. & Roe, J.-H. Induction of the sufA operon encoding Fe-S assembly proteins by superoxide generators and hydrogen peroxide: involvement of OxyR, IHF and an unidentified oxidant-responsive factor. Mol. Microbiol. 51, 1745–1755 (2004).

27. Rajagopalan, S. et al. Studies of IscR reveal a unique mechanism for metal-dependent regulation of DNA binding specificity. Nat. Struct. Mol. Biol. 20, 740–7 (2013).

28. Weiss, M. C. et al. The physiology and habitat of the last universal common ancestor. Nat. Microbiol. 1, 16116 (2016).

